# Experimentally manipulating forest structure to mimic management strategies: effects on deadwood fungal diversity and related ecosystem processes

**DOI:** 10.1101/2025.08.30.673211

**Authors:** Bronwyn Lira Dyson, Vendula Brabcová, Petr Baldrian, Jörg Müller, Michael Junginger, Claus Bässler

## Abstract

Understanding the relationships between forest management, biodiversity, and ecosystem processes is necessary for achieving multifunctionality. Deadwood fungi are extremely diverse and important for carbon turnover in forests. However, how forest structure, resulting from management, affects deadwood fungal diversity and decomposition is not well known. We experimentally tested the effects of microclimate (via canopy cover) and deadwood enrichment (snags, logs, tree crowns, and habitat trees) on fungal diversity. Further, we assessed the effects of the treatments and fungal diversity on wood mass loss. We characterized the fungal communities of *Fagus sylvatica* (European beech) and *Pinus sylvestris* (Scots pine) deadwood via metabarcoding and measured wood mass loss after 3 years. We found that the host tree species was more important than canopy cover or deadwood enrichment for fungal alpha and beta diversity. Fungal alpha diversity of beech was mainly related to canopy cover; diversity of beech was higher under closed canopies. While alpha diversity of pine was related only to deadwood enrichment as diversity increased where habitat trees and crowns remained. Furthermore, while mass loss of beech was significantly higher in patches where trees were removed and patches where crowns remained, pine mass loss was neither affected by canopy cover nor deadwood enrichment. Host tree diversity is more important than environmental variability as a determinant of fungal diversity, underpinning the importance of maintaining diverse hosts in forests. However, contrasting diversity and decomposition effects between beech and pine suggest the need for forest management strategies tailored to tree species to maintain fungal diversity and ecosystem processes.

## 1. Introduction

Modern forestry aims for multifunctionality but is faced with substantial trade-offs. One of the most important aims is to maximize wood production while maintaining biodiversity (FAO, 2024; Borrass et al., 2017). Biodiversity in forests is related to important ecosystem processes; as Liang et al. (2016) show, loss in biodiversity is linked to declines in forest productivity. Amyntas et al. (2023) connect gains in ecosystem functioning with increased niche complementarity in more diverse plant communities. To improve evidence-based forest management decisions, we require a better understanding of the complex relationships between forest management strategies and their consequences on forest structure, on the one side, and biodiversity and related ecosystem processes, on the other side.

The two key structural axes in temperate forest ecosystems that drive forest biodiversity many organism groups and related processes are microclimate (Sanczuk et al., 2023; Kemppinen et al., 2024) and deadwood of different kinds as a critical resource and habitat (Thorn et al., 2020; Stokland et al., 2012); Variations in microclimate and deadwood availability provide different abiotic constraints, opportunities, and refugia. First, many forest fungi, including threatened species, depend on deadwood as a resource (Seibold et al., 2015; Virnes et al., 2012). Clearly, a critical amount of a specific deadwood niche is necessary to maintain a vital population of a given dependent species. However, deadwood enrichment as a concept in multifunctional forestry is not straightforward; a myriad of different niches exists, due to deadwood variability (Fujii et al., 2023; Bujocwek et al., 2021). Summarized across studies, the host identity (e.g., tree species), type (e.g., logs, snags, stumps), dimension (e.g., Coarse Woody Debris, CWD, and Fine Woody Debris, FWD) and succession (decomposition stage) have been identified as the overall main drivers of saproxylic diversity (e.g., Boddy, 2021; Yang et al., 2021; Ódor et al., 2006). Second, still living but senescent (habitat) trees have been identified as key components for a broad range of rare and threatened plant and animal (Bütler et al., 2013), and fungal forest species (Siitonen, 2001). However, due to forest management intensity over centuries, these trees in forests, and the species depending on them, have become quite rare (Jones et al., 2018; Zapponi et al., 2017). This issue is well-known and thus, forest conservation biology developed the approach of inducing premature senescence (Thorn et al., 2020; Vitková et al., 2018). In this concept, mature trees still decades far from natural senescence are actively damaged in commercial forests (e.g., with a chain saw or harvester) to create habitat features analogous to those of senescent trees. Third, numerous studies have provided evidence on the importance of canopy-mediated microclimate variability for forest biodiversity (e.g., Nadkarni et al., 2001; Nakamura et al., 2017). With an opening of the canopy, the abiotic microclimate conditions change considerably leading to on average higher temperatures (Thom et al., 2020; Kovács et al., 2020) but also to variation in temperature (De Frenne et al., 2019; Braziunas et al., 2025). For example, in a recent study it has been shown that the temperature of deadwood surfaces under open canopies can reach ca. 50°C in summer, while under closed canopies the maximum was below 30°C (Schreiber et al., 2024). Altogether, it is well known that deadwood amount and variability as well as microclimate conditions are important predictors for forest biodiversity and they are directly affected by forest management. To better inform forest management and address the trade-off between timber extraction and the maintenance of forest biodiversity, we need results from experimental set-ups. These experiments can overcome confounding effects and provide evidence on the relative importance of factors driving forest biodiversity.

Multifunctional forestry not only aims to balance wood resource use with the support of forest biodiversity, but also seeks to maintain key ecosystem processes, such as primary production and the decomposition of various kinds of organic matter (Gustafsson et al., 2012; Lier et al., 2022). Particularly the latter function is related to carbon sequestration and nutrient turnover (e.g., Stokland et al., 2012; Swift 1977). The relationship between forest structure, due to forest management decisions, and ecosystem processes might be direct and/or indirect. For example, the decomposition rate of deadwood in a forest stand might be directly affected by canopy openness via physical processes (e.g., radiation/photodegradation, Fravolini et al., 2016; Gora et al., 2019). Indirectly, a change in canopy-mediated microclimate might affect decomposer communities (Gora et al., 2019; Perreault et al., 2020) with consequences on decomposition rates. However, our understanding of how forest structure affects ecosystem processes directly and indirectly via diversity change is limited (Runnel et al., 2025). Here, we address decomposition as one such ecosystem process.

We used a real-world forest experiment in which we orthogonally manipulated at the stand-level canopy cover and deadwood resources in patches of 50 m by 50 m. The independent structural deadwood features were: logs, snags, logs and snags together, tree crowns (FWD) and habitat trees. We further considered total tree removal (including stump and roots) as a treatment. As a control, we used patches in which the trees were removed but stumps were left. This reflects a continuous cover forestry regime that leaves the canopy closed, which is a standard forest management strategy in central and south-eastern Europe, Mason et al., 2022).

We manipulated ca. 30% of the living tree biomass (Kacic et al., 2024). As a further independent abiotic feature, we manipulated the canopy cover (i.e., “closed” versus “open” canopy) inducing differences in microclimate conditions. We used wood-inhabiting fungi as a model system, as this group is extremely diverse (Paterson et al., 2023; Wu et al., 2019), relevant to forest conservation (Müller et al., 2007), some are strongly dependent on old trees and deadwood (Yang et al., 2021; Runnel & Lõhmus, 2017) and, further, because fungi are the key agents driving deadwood decomposition in temperate forest ecosystems (Tláskal et al., 2021).

We sampled the microbial community of standardized exposed deadwood objects of European beech (*Fagus sylvatica* L.) and Scots pine (*Pinus sylvestris* L.) via metabarcoding and determined mass loss between 2018 and 2021. Including a broad-leaf and conifer host wood species in the design allowed us to characterize patterns in microbial fungal diversity beyond a host specific response (Jaroszewicz et al., 2021). Furthermore, *F. sylvatica* and *P. sylvestris* are two of the most common tree species in German production forests (BMEL, 2024). First, we hypothesize that the relative importance of deadwood enrichment, as habitat and food for local fungi, will be greater than that of canopy cover for explaining alpha and beta diversity. Further, diversity should increase with enrichment of CWD (i.e., logs, snags, habitat trees) and FWD (i.e., crowns) relative to the control (stumps), due to increased resource amount and variability. Consequently, fungal diversity should be lower in the total tree removal treatment than in the control. Second, we hypothesize that both canopy cover and fungal diversity will be relatively strong predictors of wood mass loss in comparison to deadwood enrichment.

## 2. Material and Methods

### 2.1 Study area and design

The study took place in the forest of the Julius-Maximilians-University of Würzburg (hereafter, “forest”) in Bavaria, Germany (Fig. 1). The forest (50.062° N, 10.444° E) is approximately 2200 ha, is an intensively managed broad-leaf forest (Kacic et al., 2024) and is mainly characterized by *Fagus sylvatica*, *Quercus* spp., *Carpinus betulus*, *Picea abies, Pinus sylvestris* and other broad-leaf (e.g., *Acer spp.)* and coniferous *(e.g., Larix ssp.)* tree species (Pierick & Ammer, 2025). The average age of the stands was between 80 and 98 years. The forest has a mean annual temperature of 8.5°C, annual precipitation of 670 mm (Stark et al., 2023), and an average elevation of 336 m.

**Figure 1.**
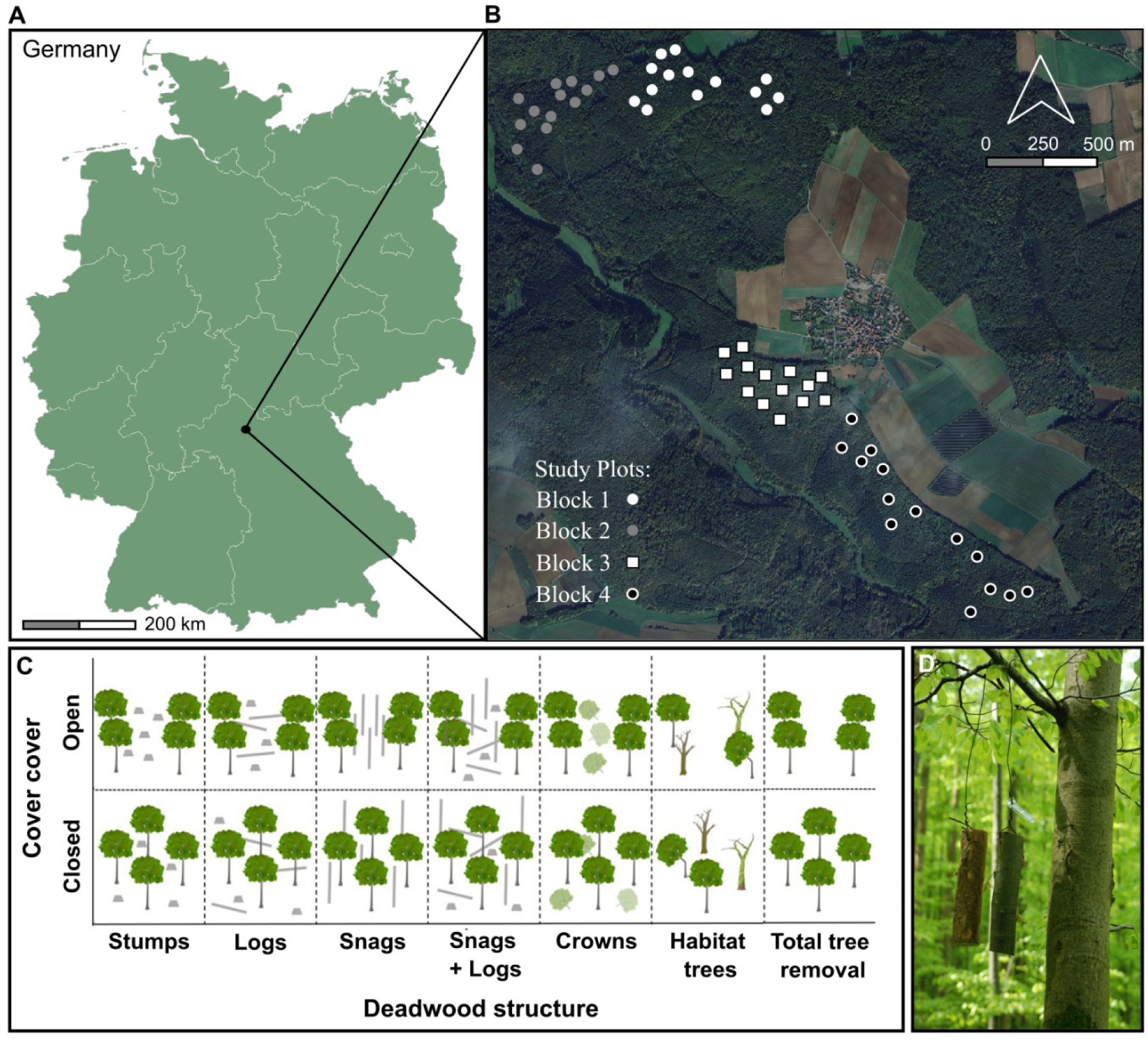
**A:** Map of Germany with study location indicated. **B:** A satellite map (© Google 2025) depicting part of the forest belonging to the Julius-Maximilians-University of Würzburg including the areas in which the study patches are located in their respective blocks as well as the town of Sailershausen. **C:** Modified from Schwarz et al., 2025. Illustration of treatments, where the top row of the illustration represents patches where the canopy is relatively open, due to the aggregated deadwood manipulation, and the bottom row represents patches where the canopy is relatively closed, due to the distributed deadwood manipulation. The control in our analyses was where stumps remain. Patches were 50 m by 50 m. **D:** A photo showing two pieces of standardized hanging wood per patch (*P. sylvestris* and *F. sylvatica*).

To address how the microclimate, through variation in canopy cover, as well as the type of deadwood available at the patch-level (i.e., log, snag, stump, habitat tree, tree crown) influenced the diversity of microbial fungi as well as deadwood mass loss, we set up an experiment in 2018 in a randomized block design through manipulating forest stands in patches (*n* = 56) of 50 m by 50 m by removing ca. 30% of the patch-level tree biomass (Kacic et al., 2024) (Fig. 1). The same approximate amounts of deadwood per patch were created irrespective of treatment type. The study patches were established across four forest blocks, hereafter “block”.

The deadwood treatments were as follows: 1) we left only stumps remaining (reference/control for our analyses) (*n* = 20), 2) we left only logs and their associated stumps (*n* = 6), 3) we left only snags (trees cut below the crown) (*n* = 6), 4) we left logs and their associated stumps, and snags (*n* = 6), 5) we left damaged trees (hereafter, “habitat” trees, i.e., with bark loss, damaged crowns, or tree hollows; Müller et al., 2014) (*n* = 6), 6) we removed trees but left tree crowns (other than in this treatment, tree crowns were always removed) (*n* = 6), and 7) trees, including stumps, were completely removed from the plots (*n* = 6). The reference treatment (stumps) best resembles the dominant tree harvesting practice in central European forests (Mason et al., 2022).

For the microclimate aspect of the treatment, we had: 1) aggregated deadwood manipulation, where we created canopy gaps by cutting trees in groups of approximately 625 m² (*n* = 20) and 2) distributed deadwood manipulation, where we avoided creating large canopy gaps by cutting trees spaced apart from one another across the entire 50 m by 50 m patch (*n* = 36). The individual tree cuts permit relatively less light to penetrate through the canopy compared to the aggregated tree cuts. Closed canopy treatments resembled a single-tree selection forest management strategy, while the open canopy treatments resembled rather group-selection cuts (Schall et al., 2017, Goßner et al., 2006) or a small-scale natural disturbance like local windthrow (Forrester et al., 2012). For simplicity, we refer to the canopy treatments as “open” and “closed” hereafter.

On each of the patches, we hung two logs of deadwood of two host trees, *Fagus sylvatica* (hereafter, “*F. sylvatica*”) and *Pinus sylvestris* (hereafter, “*P. sylvestris*”). We hung the logs from tree branches close to the patch center or, where no trees were available, on wooden posts in the patch center. We hung the logs to focus our study on the colonization of wood by airborne fungal spores, excluding soil fungi colonization. The logs were approximately 15 cm long with a diameter of 6 cm.

### 2.2 Sampling

In 2018 and in 2021 we sawed an approximately 4 cm slice from the hanging logs and froze them at -40°C. We used both slices to determine mass loss (see below) and the 2021 slice to analyze the microbial communities. For this purpose, we defrosted the slices directly before drilling a sawdust sample. We used a drill (Makita, Japan) and an 8-mm diameter drill bit for collecting the deadwood sample. The drill was cleaned and sterilized with ethanol and flame before each slice was drilled. Two perpendicular drillings per slice were taken. We collected the sawdust directly into sterile plastic bags, which we froze immediately after drilling to conserve the DNA. The sawdust samples were then freeze-dried and milled using the Ultra Centrifugal Mill ZM 200 (Retsch, Germany).

### 2.3 DNA extraction, PCR, amplicon sequencing, and OTU clustering

Approximately 200 mg of freeze-dried material was used to extract DNA via the NucleoSpin Bead Tube Type A (Macherey-Nagel, Germany). After adding Enhancer SX, we used an SL1 lysis buffer to lyse cells. We homogenized samples using FastPrep-24 (MP Biomedicals, Santa Anna, USA) at 6.5 m s -1 for 2 × 30 seconds (Baldrian et al., 2016). Finally, we eluted DNA from columns using 50 μl of deionized water. DNA quality and quantity were assessed using a NanoDrop spectrophotometer (NanoDrop 2000, Thermo Scientific, USA).

The fungal community composition was analyzed by sequencing the ITS2 region of DNA using barcoded primers gITS7 and ITS4 (Baldrian et al., 2016). To minimize PCR bias, polymerase chain reaction (PCR) was performed in triplicate for each DNA sample. Each PCR contained 5 µl of 5× Q5 reaction buffer, 1.5 µl of BSA (10 mg ml-1), 1 µl of each primer (0.01 mM), 0.5 µl of PCR Nucleotide Mix (10 mM each), 0.25 µl of Q5 High Fidelity DNA polymerase (2 U µl -1, New England Biolabs, Inc.), 5 µl of 5× Q5 HighGC Enhancer and 1 µl of template DNA (approx. 50 ng µl-1). Cycling conditions were 98°C for 30 sec, 30 cycles of 94°C for 10 sec, 56°C for 30 sec, and 72°C for 30 sec, and a final extension at 72°C for 2 min. Negative and positive amplification controls were used to avoid contamination errors and to prove the efficiency of the PCR, but these controls were not sequenced within this project. PCR triplicate reaction products were pooled and purified (MinElute PCR Purification Kit, Quiagen), and amplicon libraries prepared with the TruSeq DNA PCR-Free Kit LP (Illumina).

Fungal amplicon sequencing was conducted at the Institute of Microbiology at the Czech Academy of Sciences (Prague, Czech Republic), using an Illumina MiSeq (2 × 250 base pair-end run) (Illumina, USA) as described previously (Baldrian et al., 2016). We processed the amplicon sequencing data using the pipeline SEED 2.1.3 (Větrovský and Baldrian, 2013). We merged pair-end reads using fastq-join (Aronesty, 2013) and extracted the ITS2 region with ITS Extractor 1.0.11 (Bengtsson-Palme et al., 2013). Using Usearch 11.0.667 (Edgar, 2010), we detected chimeric sequences and removed them. Clustering of sequences into Operational Taxonomical Units (OTUs) was done using the UPARSE algorithm implemented within Usearch 8.1.1861 (Edgar, 2013) at a 97% similarity level. The most abundant sequence was used to represent each OTU.

### 2.4 Data preparation: sample reads threshold, singletons, and sample coverage

We established a minimum read sum of 2000 for analyses (i.e., all samples had sufficient reads). Then, we used the approach of Chiu & Chao (2016) to estimate the true number of singletons and we modified the raw community matrix accordingly. Specifically, this approach modifies an OTU table by randomly setting singleton entries to zero such that the number of remaining singletons per sample is equal to the number of singletons estimated by Chiu & Chao’s method for sequencing error estimation. As this procedure involves a resampling approach, multiple runs (i.e., modified community matrices) should ideally be considered in the subsequent analysis. However, testing different resampled community matrices did not alter the inference of our final models. Therefore, for simplicity and reproducibility, we used *set.seed* (R Core Team, 2022) and used a single representative community matrix after singleton correction.

Next, we calculated sample coverage to avoid sampling bias effects in our analyses. Sampling bias or sample incompleteness is an issue in ecological research which can affect a study’s inferences (Chao & Jost, 2012). For example, in our data, sampling bias can result from differences in detection probability (e.g., sequencing depth) caused by differences in environmental conditions (Kortmann et al., 2025). We calculated sample coverage using the *iNEXT* function and package (Chao et al., 2014a; Chao et al., 2014b; Hsieh et al., 2016) with datatype = “abundance”. We then continued by assessing potential biases in sample coverage in response to predictor variables using beta regression models, as sample coverage values are between 0 and 1. We used the *gam* function from the *mgcv* package (Wood, 2011), specifying plot in block as nested random effects (family = “betar”, link = “logit”). The response variable was sample coverage and the fixed effects were tree species, canopy, and deadwood treatment. Biases (i.e., statistically significant effects of predictors on sample coverage) would suggest that a standardized level of sample coverage for diversity analyses is not only appropriate but necessary. We found that tree species had significant effects on sample coverage – indicating that standardization based on sample coverage was necessary for further analyses.

### 2.5 Calculating alpha and beta diversity with the Hill numbers

The Hill numbers can be understood as the effective or true diversities of communities (Marion et al., 2021; Hill, 1973; Ellison, 2010). The adjustable parameter *q* indicates how sensitive the index is to species abundances (Chao et al., 2014). *q* = 0 corresponds to species richness, where species abundances are irrelevant, and a disproportionate weight is thus given to rare species. *q* = 1 corresponds to Shannon diversity, where species are weighted proportionate to their abundances and common species are thus emphasized. *q* = 2 corresponds to the inverse of Simpson’s diversity, where dominant species are emphasized and rare species are discounted (Chao et al., 2014). A clear advantage to using the Hill numbers framework is the possibility it offers to assess perhaps distinct responses and patterns across different abundance classes of species. For simplicity, we refer to the OTUs hereafter as “species.”

To calculate alpha diversity, for *q* = 0, 1, and 2, we used the *estimadeD* function from the *iNEXT* package (Chao et al., 2014; Hsieh et al., 2022), with the settings: datatype = “abundance”, base = “coverage”, level = NULL. When level = NULL, this is the default setting; the function then computes the diversity estimates for the minimum among all the coverage values for samples extrapolated to double the reference sample size.

To calculate beta diversity, for the Hill numbers *q* = 0, 1, and 2, we used the *iNEXTbeta3D_pair3D* function (Chao et al., 2023; Kortmann et al., 2025) with the following arguments: datatype = “abundance”, parallel = T, cpus = 12, SC = Cmax_joint. Where *q* = 0, 1, and 2 correspond to the Sørensen, Horn, and Morisita-Horn indices, respectively. The input value for sample coverage (“Cmax_joint”), was computed as the minimum among all the coverage values for samples extrapolated to double the reference sample sizes (Chao et al., 2023). We then used the beta diversity results of each Hill number to create distance matrices for the fungal community using the functions *matrix* and *as.dist* from the *base* and *stats* packages, respectively (R Core Team, 2022). We conducted a Principal Coordinates Analysis (PCoA) based on the distance matrices using the *pco* function from the *ecodist* package (Goslee & Urban, 2007). We then extracted the scores of the first and second axes from the PCoA to represent beta diversity and act as variables in our models (Kümmet et al., 2025).

### 2.6 Measuring mass loss

We dried the wood slices from 2018 and 2021 at 50°C until the mass was constant. We removed the few fruit bodies that occurred on a few slices. Next, we measured the dry mass of the slices with a fine scale (i.e., accurate to three decimal points). We then wrapped the wood slice in Parafilm (Bemis Company, USA) and determined its volume via water displacement using a glass volumetric vessel. To determine density, we divided the dry weight by the volume. Percent mass loss from 2018-2021 was calculated as follows: ((*dry weight/volume*^2018^ – *dry weight*/*volume*^2021^)/(*dry weight*/*volume*^2018^)) * 100.

### 2.7 Heteroscedasticity, response distribution, and model choice

To determine if it was appropriate to model *F. sylvatica* and *P. sylvestris* deadwood samples together or separately, we ran Levene tests (*leveneTest* function from the *car* package, Fox & Weisberg, 2019) to discern if the variance between *F. sylvatica* and *P. sylvestris* sample subsets was homogenous or not, in terms of alpha and beta diversity as well as with mass loss as model responses across orders of *q*. The results of the Levene test (with tree species as predictor of diversity/mass loss) showed that there was heteroscedasticity based on host tree species which led us to model the tree species together as well as separately. We repeated this approach to ascertain the homogeneity of variance based on canopy cover and deadwood treatments but found no evidence of heteroscedasticity.

For modelling alpha diversity in response to the treatments, we ran a negative binomial model with a log link using the *glmmTMB* function and package (Brooks et al., 2017) for both tree species samples together (hereafter “overall” model) as well as each of the tree species separately (hereafter “host tree species” model) and for each order of *q* (i.e., 0, 1, and 2). The rounded alpha diversity estimate was the response with canopy cover and deadwood treatment as fixed effects and with patch nested in block as random effects for the overall models and block as a random effect in the host tree species specific models. Additionally, for *q* = 0, since there was heteroscedasticity in terms of the host tree species, we ran a robust linear mixed-effects model using the *rlmer* function from the *robustlmm* package (Koller, 2016) to verify that the results from the *glmmTMB* were robust.

For modelling beta diversity, we used the scores of the first and second axes (PC1 and PC2, respectively) from the PCoA as response variables in an overall model as well as host tree species models. We included canopy cover and deadwood treatment as fixed effects with patch nested in block as random effects for the overall models and block as a random effect in the host tree species specific models. We used Linear Mixed Effects models with *lmer* function from the *lme4* package (Bates et al., 2015). Using the scores from the PCoA axes as model responses has the advantage of permitting an analysis that fits in the same statistical framework as the alpha diversity analyses. Importantly, it allows for testing the levels of our factors in relation to a reference (i.e., the control patches) as in the alpha models. However, based on this approach, we do not use the full variance of the ordination – although it is important to note that we explain considerable variability with the first two axes alone (across tree species and orders of *q*). Applying additional models with up to four PCoA axes, applying a PERMANOVA, or calculating environmental fits directly on to the ordination (function *envfit*, *vegan* package; Oksanen et al., 2025) yielded no further insights (data not shown).

To assess simultaneously direct and indirect effects of canopy cover, deadwood enrichment, and host tree species on mass loss as well as the effects on alpha and beta diversity in an overall model, we used a Structural Equation Model (SEM) from the *psem* function from the *piecewiseSEM* package (Lefcheck, 2016). We followed the approach put forth by Lefcheck (2021) for running SEMs with exogenous categorical variables, whereby a built-in ANOVA tests for the effect of the categorical variable and a follow-up test assesses the model-estimated or marginal means for each group level. We square root transformed mass loss and log-transformed alpha diversity to achieve normal distributions. We used linear mixed effects (LME) models within the SEMs to account for patch in block as nested random effects (method = “ML”). For each order of *q* we used two LME models within the SEM. The first LME was a “diversity as response” model: diversity was the response to canopy cover and deadwood treatment. The second LME was “mass loss as a response” model: the mass loss proportion was the response to canopy cover, deadwood treatment, and diversity. We ran three separate SEMs for each diversity variable (i.e. alpha diversity, beta diversity (PC1), and beta diversity (PC2)). Note that, to account for the different diversity estimates (alpha and beta diversity across orders of *q*) used as predictors in the SEMs, we fitted for each diversity measure one separate SEM. As the other predictors (i.e., canopy and deadwood treatments as well as host tree species) were kept constant, we only consider these predictors significant when they had significant effects across all models (i.e., for each order of *q* and diversity estimate).

For host tree species specific models, we used beta regressions for assessing the effect of canopy cover, deadwood treatment, and diversity on the proportion of mass loss in each of the host tree species separately and for each diversity estimate separately (i.e., alpha diversity, beta diversity (PC1), and beta diversity (PC2)). We log-transformed alpha diversity. We used the *gam* function from *mgcv* package (Wood, 2011) with family = “betar” and link = “logit” and we included block as a random effect. We report deviance explained as the models are from a non-Gaussian family and thus have non-normal errors (Wood, 2011; Clark, 2022). Furthermore, we confirmed that there were no issues of multicollinearity between predictors in the models by using the *vif* function from the *car* package (Fox & Weisberg, 2019).

## 3. Results

### 3.1 Effects of canopy cover, deadwood treatments, and host tree species on fungal alpha diversity

#### Overall model

Among all predictors, host tree species showed the strongest effect on alpha diversity for all orders of *q* (Table 1). Alpha diversity was consistently higher on *P. sylvestris* than on *F. sylvatica*. The relative importance of the deadwood treatment and the canopy cover treatment on alpha diversity differed among the orders of *q*. While for the diversity of rare species the effect of canopy cover was significant, the deadwood treatments were not significant for any abundance class of species, although the habitat trees treatment was near significant for common species and the crowns treatment was near significant for dominant species (Table 1). However, the observed overall effects were driven by host tree species-specific responses.

**Table 1.** Test statistics for fungal alpha diversity using negative binomial models with a log link. In the overall model, *F. sylvatica* and *P. sylvestris* samples were modelled together. Alpha diversity estimates were rounded before modelling. Patch nested in block were random effects for the overall model, block was a random effect in the host tree species specific models. Reference levels for canopy and host tree are indicated in parentheses. The reference level for deadwood treatment was stumps, i.e., patches where only stumps were remaining. Significant effects are indicated as: * *p* < 0.05, ** *p* < 0.01, *** *p* < 0.001.

| | Fixed effects | $q = 0$, rare | | $q = 1$, common | | $q = 2$, dominant | |
| --- | --- | --- | --- | --- | --- | --- | --- |
|  |  | z value | Pr(>\|z\|) | z value | Pr(>\|z\|) | z value | Pr(>\|z\|) |
| Overall model | Canopy (Open) | **2.740** | **0.006\*\*** | 1.493 | 0.136 | 1.482 | 0.138 |
|  | Trees removed | 1.002 | 0.316 | 0.548 | 0.584 | 0.412 | 0.681 |
|  | Snags + Logs | -0.082 | 0.935 | -0.342 | 0.732 | -0.665 | 0.506 |
|  | Habitat trees | 1.416 | 0.157 | 1.905 | 0.057 | 1.404 | 0.160 |
|  | Crowns | 0.879 | 0.379 | 1.326 | 0.185 | 1.952 | 0.051 |
|  | Logs | -0.492 | 0.623 | -0.648 | 0.517 | -0.985 | 0.324 |
|  | Snags | -0.032 | 0.975 | 0.329 | 0.742 | 0.305 | 0.760 |
|  | Host tree (*F. sylvatica*) | **9.224** | **<0.001\*\*\*** | **6.338** | **<0.001\*\*\*** | **3.963** | **<0.001\*\*\*** |
|  | Marginal R<sup>2</sup> | 0.482 |  | 0.325 |  | 0.213 |  |
| *F. sylvatica* | Canopy (Open) | **3.192** | **0.001\*\*** | **2.734** | **0.006\*\*** | **2.248** | **0.025\*** |
|  | Trees removed | 1.524 | 0.128 | 0.922 | 0.357 | 0.889 | 0.374 |
|  | Snags + Logs | 0.689 | 0.491 | 0.766 | 0.444 | 0.605 | 0.545 |
|  | Habitat trees | 0.631 | 0.528 | 0.580 | 0.562 | 0.425 | 0.671 |
|  | Crowns | 0.869 | 0.385 | 0.275 | 0.783 | 0.444 | 0.657 |
|  | Logs | -0.781 | 0.435 | -0.506 | 0.613 | -0.826 | 0.409 |
|  | Snags | 0.922 | 0.356 | 0.997 | 0.319 | 0.597 | 0.550 |
|  | Marginal R<sup>2</sup> | 0.222 |  | 0.169 |  | 0.141 |  |
| *P. sylvestris* | Canopy (Open) | 0.265 | 0.791 | -0.710 | 0.477 | 0.005 | 0.996 |
|  | Trees removed | 0.650 | 0.516 | 0.470 | 0.638 | 0.012 | 0.990 |
|  | Snags + Logs | 0.018 | 0.985 | -0.637 | 0.524 | -1.243 | 0.214 |
|  | Habitat trees | **2.098** | **0.036 \*** | **2.659** | **0.008\*\*** | 1.671 | 0.095 |
|  | Crowns | 1.273 | 0.203 | **2.236** | **0.025\*** | **2.358** | **0.018\*** |
|  | Logs | 0.549 | 0.583 | 0.027 | 0.978 | -0.473 | 0.637 |
|  | Snags | -0.591 | 0.554 | 0.030 | 0.976 | 0.129 | 0.898 |
|  | Marginal R<sup>2</sup> | 0.138 |  | 0.218 |  | 0.198 |  |

#### Host tree species specific models

When modelling the response of alpha diversity for each of the host tree species separately, for *F. sylvatica*, fungal diversity was consistently (i.e., for *q* = 0, 1, and 2) higher in patches where the canopy was closed (Fig. 2). The fungal diversity of *P. sylvestris* was higher in patches where habitat trees were present, in terms of rare and common species, and where and tree crowns were present, in terms of common and dominant species (Fig. 2).

**Figure 2.**
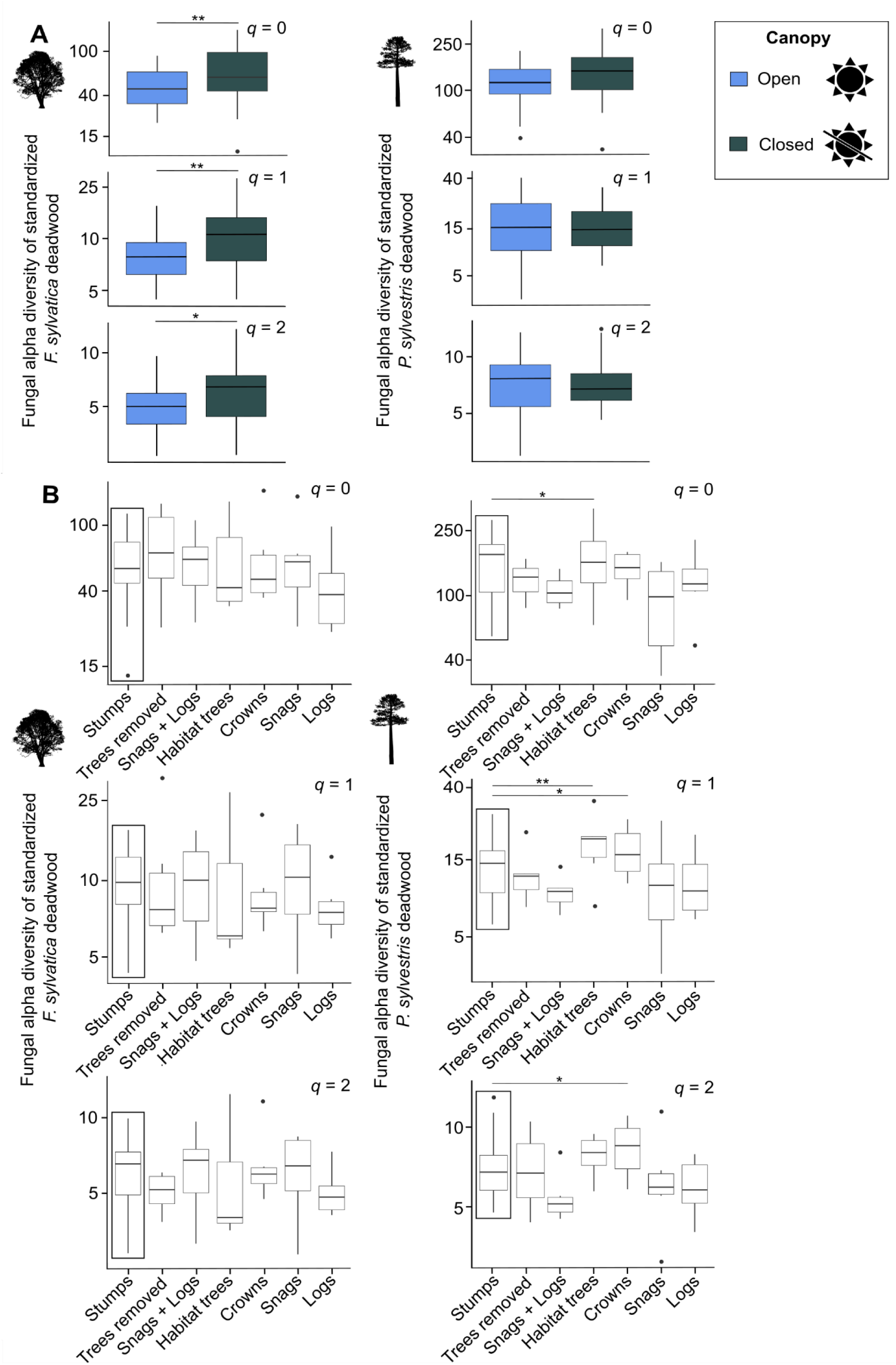
Fungal alpha diversity across orders of *q* (i.e., 0 = rare species, 1 = common species, 2 = dominant species) for *F. sylvatica* and *P. sylvestris* host tree standardized deadwood in response to the canopy cover (**A**) and deadwood treatments (**B**). Diversity values were log-transformed. The control for the comparisons among deadwood were patches with stumps remaining (boxed in black). Contrasts to the control that were significant from the host tree species specific models (see Table 1 for complete results) are indicated with asterisks (* *p* < 0.05, ** *p* < 0.01, *** *p* < 0.001).

### 3.2 Effects of canopy cover, deadwood treatments, and host tree species on fungal beta diversity

#### Overall model

Similar to fungal alpha diversity, fungal beta diversity was mainly explained by host tree species across all orders of *q* (Table 2). The canopy treatment was more important than the deadwood enrichment treatments, specifically, the beta diversity of rare species was significantly related canopy (Table 2). We found no significant effect of deadwood treatments for beta diversity.

**Table 2.** Test statistics from fitting linear mixed effects models to the scores of the first and second axes (PCs) of the PCoA for the overall model and for *F. sylvatica* and *P. sylvestris* specific models in response to canopy, deadwood treatments, and (in the case of the overall model) host tree species. Reference levels for canopy and host tree are indicated in parentheses. The reference level for deadwood treatment was stumps, i.e., patches where only stumps were left remaining. Significant effects are indicated as: * *p* < 0.05, ** *p* < 0.01, *** *p* < 0.001.

| Fixed effects |  | *q* = 0, rare |  |  |  | *q* = 1, common |  |  |  | *q* = 2, dominant |  |  |  |
| --- | --- | --- | --- | --- | --- | --- | --- | --- | --- | --- | --- | --- | --- |
|  |  | PC1 |  | PC2 |  | PC1 |  | PC2 |  | PC1 |  | PC2 |  |
|  |  | t value | Pr(>\|t\|) | t value | Pr(>\|t\|) | t value | Pr(>\|t\|) | t value | Pr(>\|t\|) | t value | Pr(>\|t\|) | t value | Pr(>\|t\|) |
| Overall model | Canopy (Open) | 1.760 | 0.082 | **-2.501** | **0.016\*\*** | 1.548 | 0.125 | -0.735 | 0.464 | 1.473 | 0.147 | 1.282 | 0.203 |
|  | Trees removed | 1.537 | 0.127 | -0.266 | 0.792 | 0.644 | 0.522 | 0.391 | 0.697 | 0.364 | 0.718 | -0.444 | 0.659 |
|  | Snags + Logs | -1.446 | 0.151 | -1.555 | 0.127 | 0.349 | 0.728 | -1.432 | 0.157 | 0.560 | 0.579 | 0.905 | 0.369 |
|  | Habitat trees | -1.357 | 0.178 | -1.789 | 0.080 | 0.524 | 0.603 | -0.208 | 0.836 | 0.445 | 0.659 | -0.541 | 0.590 |
|  | Crowns | -1.575 | 0.118 | -1.11 | 0.272 | -0.501 | 0.619 | -0.793 | 0.431 | -0.060 | 0.953 | 0.276 | 0.783 |
|  | Logs | -1.755 | 0.082 | -0.016 | 0.987 | -0.424 | 0.673 | 0.919 | 0.361 | -0.465 | 0.645 | -1.441 | 0.154 |
|  | Snags | -0.692 | 0.491 | -0.311 | 0.757 | 0.687 | 0.495 | 0.001 | 0.999 | 1.191 | 0.241 | -0.190 | 0.850 |
|  | Host tree (*F. sylvatica*) | **45.190** | **<0.001\***<br>**\*\*** | 0.862 | 0.393 | **45.789** | **<0.001\***<br>**\*\*** | 0.654 | 0.514 | **31.955** | **<0.001\***<br>**\*\*** | -1.068 | 0.288 |
|  | Marginal R<sup>2</sup> | 0.949 |  | 0.096 |  | 0.950 |  | 0.057 |  | 0.896 |  | 0.073 |  |
| *F. sylvatica* | Canopy (Open) | **-2.237** | **0.030\*** | **-2.206** | **0.032\*** | -1.419 | 0.162 | 0.302 | 0.764 | -1.575 | 0.122 | -0.110 | 0.913 |
|  | Trees removed | -1.089 | 0.282 | 0.766 | 0.448 | 0.101 | 0.920 | 1.403 | 0.168 | 0.196 | 0.846 | 1.466 | 0.151 |
|  | Snags + Logs | -1.116 | 0.271 | 0.372 | 0.712 | -1.568 | 0.124 | -1.385 | 0.174 | -0.803 | 0.427 | -1.227 | 0.227 |
|  | Habitat trees | -0.432 | 0.668 | 0.222 | 0.826 | -0.088 | 0.930 | -1.310 | 0.198 | 0.582 | 0.564 | -0.738 | 0.465 |
|  | Crowns | -0.874 | 0.387 | 0.694 | 0.491 | -0.807 | 0.424 | -1.323 | 0.193 | -0.142 | 0.887 | -1.653 | 0.106 |
|  | Logs | 0.603 | 0.550 | 0.915 | 0.365 | 0.848 | 0.401 | -1.261 | 0.215 | 1.638 | 0.110 | -0.941 | 0.352 |
|  | Snags | -0.592 | 0.557 | 1.590 | 0.119 | -0.085 | 0.932 | 0.026 | 0.980 | -0.083 | 0.934 | 0.344 | 0.732 |
|  | Marginal R<sup>2</sup> | 0.119 |  | 0.146 |  | 0.108 |  | 0.165 |  | 0.129 |  | 0.156 |  |
| *P. sylvestris* | Canopy (Open) | 0.166 | 0.869 | 0.016 | 0.987 | 1.837 | 0.073 | -1.739 | 0.089 | 1.803 | 0.078 | 1.726 | 0.091 |
|  | Trees removed | 1.206 | 0.234 | -1.513 | 0.137 | -0.148 | 0.883 | 0.169 | 0.866 | 0.282 | 0.779 | 0.391 | 0.697 |
|  | Snags + Logs | -0.151 | 0.881 | 0.340 | 0.735 | -0.257 | 0.799 | -0.186 | 0.853 | 0.237 | 0.813 | 0.440 | 0.662 |
|  | Habitat trees | 0.630 | 0.532 | -0.453 | 0.653 | -1.186 | 0.242 | -0.787 | 0.435 | -0.881 | 0.383 | 1.252 | 0.217 |
|  | Crowns | **2.079** | **0.043\*** | 0.714 | 0.479 | -1.951 | 0.057 | 0.367 | 0.715 | -1.702 | 0.095 | 0.235 | 0.815 |
|  | Logs | 0.959 | 0.343 | 0.811 | 0.422 | -0.663 | 0.511 | -0.399 | 0.692 | -0.154 | 0.879 | 0.781 | 0.439 |
|  | Snags | 0.348 | 0.729 | 0.564 | 0.575 | -1.192 | 0.239 | -0.466 | 0.643 | -1.126 | 0.266 | 0.655 | 0.516 |
|  | Marginal R<sup>2</sup> | 0.044 |  | 0.094 |  | 0.112 |  | 0.057 |  | 0.112 |  | 0.057 |  |

#### Host tree species-specific models

We found significant effects of canopy cover on the beta diversity of rare species of *F. sylvatica* (Fig. 3, Table 2). The amount of variance explained for PC1 of *F. sylvatica* was approx. 12-30% and for PC2 7-27% (Fig. 3). The fungal beta diversity of *P. sylvestris* was only affected by the deadwood treatment with crowns (Fig. 3, Table 2). The range in variance explained for PC1 of *P. sylvestris* was approx. 13-25% and for PC2 9-20% (Fig. 3).

**Figure 3.**
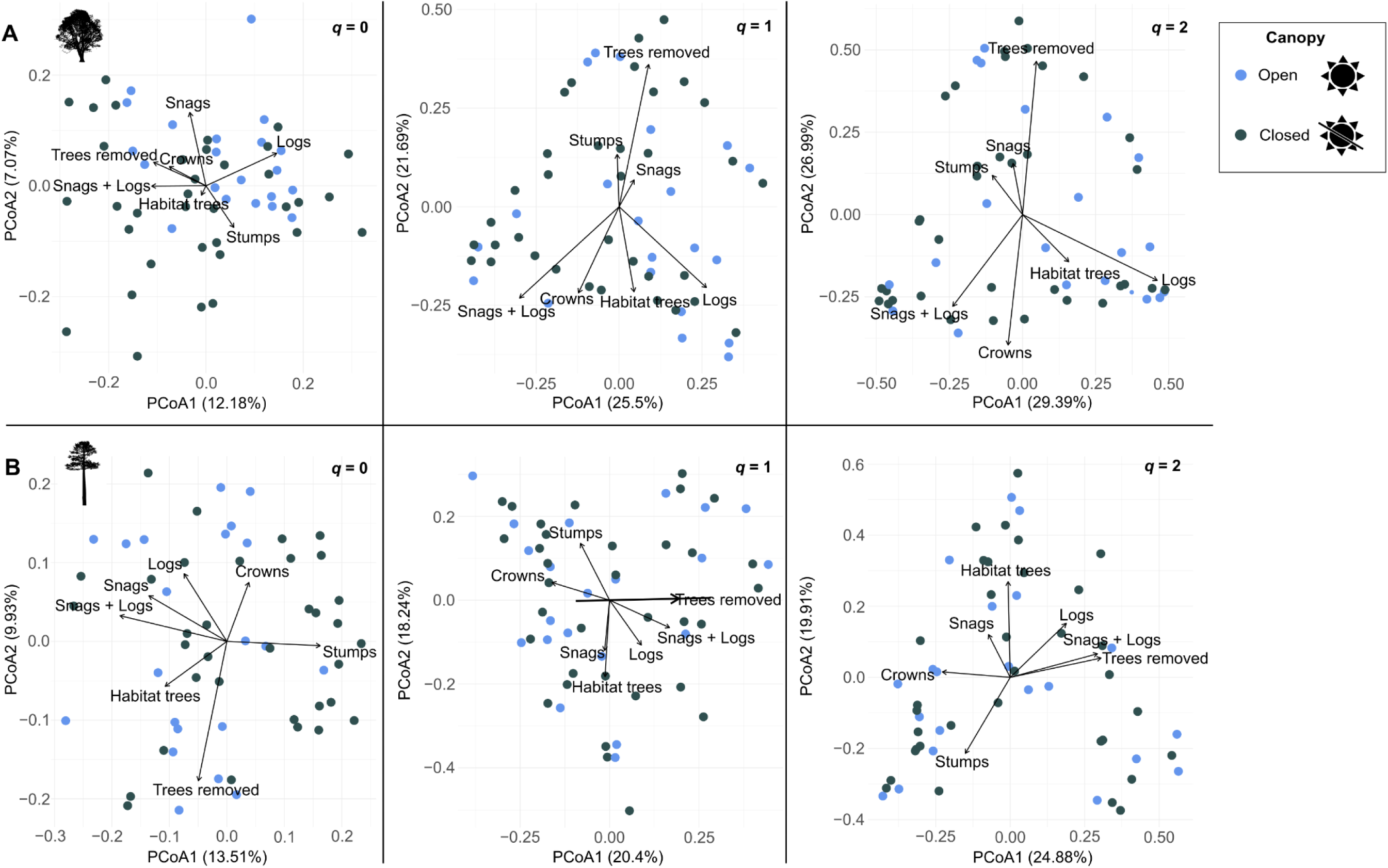
Ordination plots of the scores of the first and second axes of the PCoA conducted on the distance matrices of the pairwise *β*-diversity indices that we computed for *F. sylvatica* (**A**) and *P. sylvestris* (**B**) with canopy (colored points) and deadwood (arrows) treatments indicated along the orders of *q*.

### 3.3 Effects of canopy, deadwood treatments, and host tree species on decomposition

#### Overall Piecewise Structural Equation Model (SEM)

In an overall SEM, we analyzed the *F. sylvatica* and *P. sylvestris* samples together to assess the effects of canopy cover, deadwood treatment and host tree species on fungal diversity and mass loss. The models revealed that the environmental variables had no direct significant effect on deadwood mass loss (Fig. 4, Table S1). In contrast, alpha diversity of common and dominant species was significantly negatively related to mass loss (Fig. 4, Table S1). Further, we found that the host tree significantly affected alpha and beta diversity (Fig. 4, Table S1) which is in line with the explicit diversity models above (sections 3.1 and 3.2).

**Figure 4.**
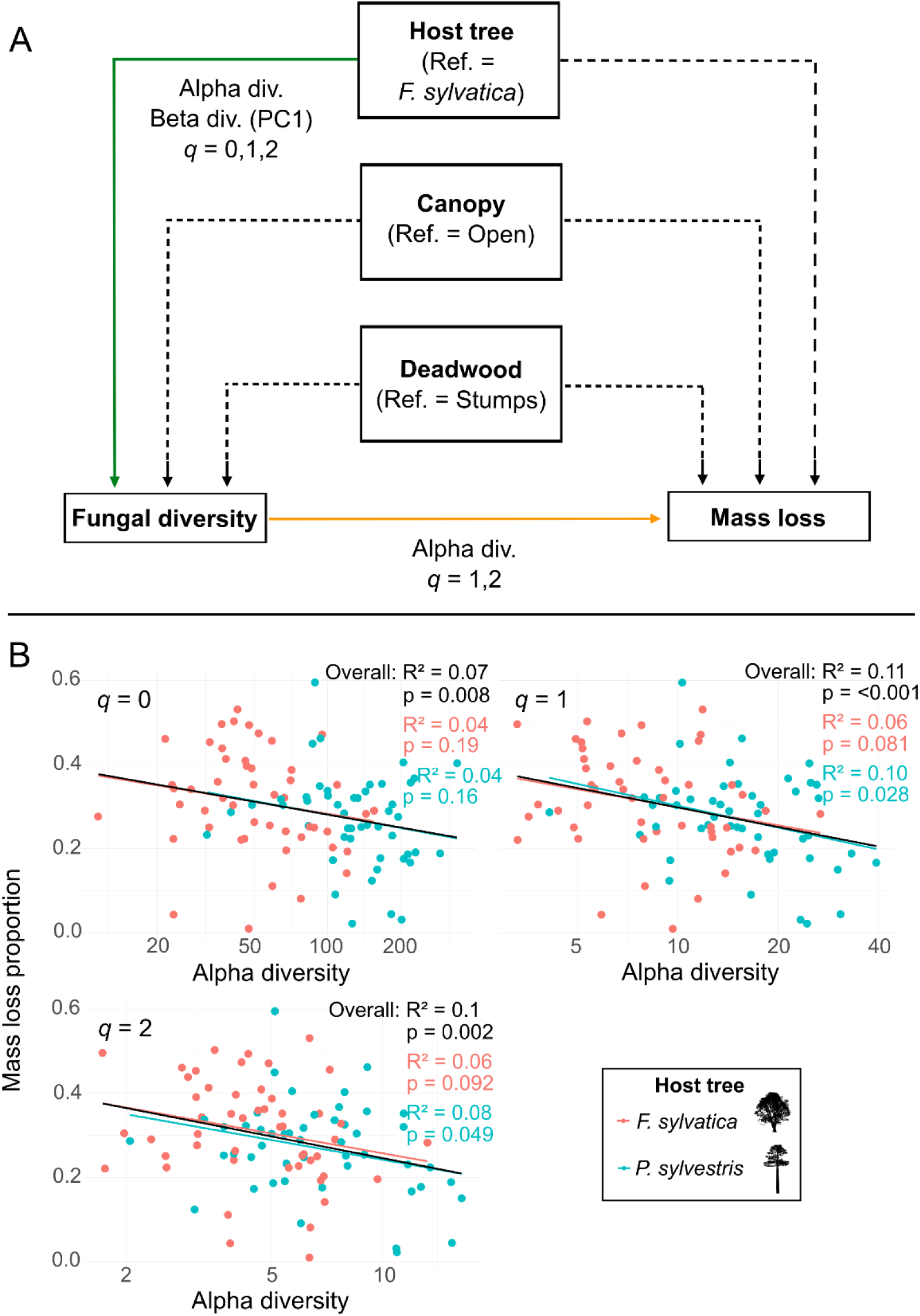
**A:** Summarized results from the overall Structural Equation Models (SEM). Reference levels for categorical predictors are stated. Orange bold line indicates significant negative relationship, green bold line indicates significant positive relationship, e.g., *P. sylvestris* had significantly higher (green line) diversity than *F. sylvatica*. Dashed lines indicate no significant relationship. In the case of fungal diversity, the relevant diversity estimate is indicated. **B:** Linear regressions of mass loss proportion (square root transformed) as a function of alpha diversity (log-transformed; values on *x* axis have been back-transformed for ease of interpretation) to illustrate the diversity-mass loss relationships tested for in the overall SEM. The linear regression R^2^ values are based on a univariate test of the diversity-mass loss relationship using *lm* from the *stats* package (R Core Team, 2022).

#### Host tree species specific beta regression models

Similar to our approach for assessing significance in the SEMs, we only considered predictors significant when having significant effects across all beta regression models (i.e., across orders of *q* and diversity estimates). In the *F. sylvatica* models, neither canopy, alpha, nor beta diversity had significant effects on mass loss. However, across orders of *q* and including models of each diversity estimate, deadwood in patches with trees removed or with tree crowns exhibited significantly higher mass loss compared to deadwood in control patches where only stumps remained (Table S2, Fig. 5). In two of the three *P. sylvestris* models (*q* = 1, 2) which included PC2, we found a significant relationship between beta diversity and mass loss (Table S2).

**Figure 5.**
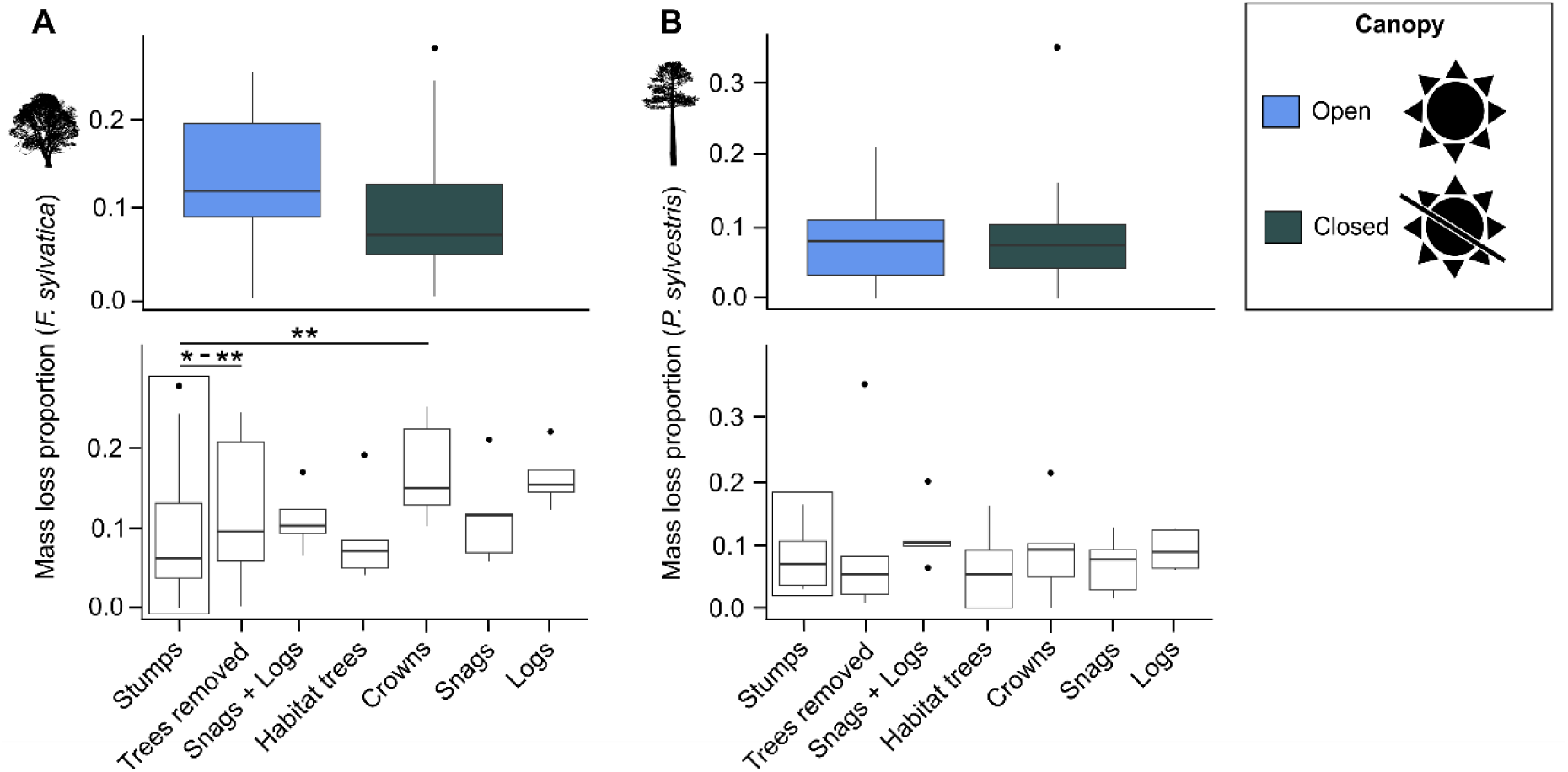
Proportional mass loss after three years for *F. sylvatica* (**A**) and *P. sylvestris* (**B**) in response to canopy and deadwood treatment. The control for the comparisons among deadwood was patches with stumps remaining (boxed in black). Contrasts to the control that were significant are indicated with asterisks (* *p* < 0.05, ** *p* < 0.01, *** *p* < 0.001); note, since nine models were run per host tree species, significance is depicted as a range where models resulted in different *p* values.

## 4. Discussion

Our large-scale experiment revealed that the host tree identity was a better predictor of fungal diversity in deadwood after three years of decomposition than forest structure-related variables. Canopy cover and deadwood resources significantly affected fungal diversity but the relative importance of canopy cover and deadwood for explaining fungal diversity differed between host tree species. Deadwood mass loss was affected by both the deadwood treatments and fungal diversity, however, the response was tree-species specific.

### 4.1 Effects of canopy cover, deadwood treatments, and host tree species on fungal alpha and beta diversity

Our overall models revealed that host tree species is more important for explaining fungal diversity than variability in environmental variables which, in our study, were represented by canopy cover and deadwood. This finding is in line with previous studies demonstrating the importance of the host identity relative to abiotic factors. As these studies were based on metabarcoding (Englmeier et al., 2023; Brabcová et al., 2022) and fruit bodies (Krah et al., 2018a; Baber et al., 2016) as well as in and across different macroclimate regimes (Rieker et al 2022), we conclude that this is a general phenomenon at the scale within and across landscapes in a biogeographic region. One explanation might be that wood-inhabiting fungi share an intimate long-lasting evolutionary relationship with their hosts and coevolved across a broad range of environmental conditions in which the hosts could survive (Floudas et al., 2012; Krah et al., 2018b). However, it seems less clear whether broad-leaf or coniferous trees host more diverse wood-inhabiting fungal communities. Our study revealed that *P. sylvestris* is more diverse in species than *F. sylvatica*. This is line with Purahong et al. (2018), who showed that coniferous logs hosted higher fungal alpha and gamma diversity than broad-leaf logs. Similarly, and more relevant to our study as FWD were analyzed, Brabcová et al. (2022) found higher fungal alpha in *Abies alba* than in *F. sylvatica* FWD. However, such findings contrast with Krah et al. (2018a), in which the authors compared the diversity between *F. sylvatica* and *A. alba* and found that *F. sylvatica* hosted higher fungal diversity than *A. alba*. One important difference in Krah et al. (2018a) is that the authors used fruit body inventories, whereas in our study, in Purahong et al. (2018), and in Brabcová et al. (2022), metabarcoding data was used. Thus, the prevalence of fruiting cues may differ between broad-leaf and coniferous trees and this may explain the contrasting results based on fruit body surveys versus metabarcoding (Rieker et al., 2024; Blaschke et al., 2023). On the other hand, generalizing about diversity patterns simply based on the tree being an angio- or gymnosperm can be a misleading contrast, as the patterns may also be more specifically related to individual tree species. One further issue is that most comparisons in the literature are based on experiments in which both broad-leaf and coniferous deadwood objects have been exposed at the same time. Comparing the fungal diversity among tree species after a defined time (e.g., after 3 years) might simply reflect differences in successional (i.e., decomposition) stages. From previous studies we know that deadwood succession is correlated with species richness (Schreiber et al 2024; Rajala et al., 2010). By measuring mass loss in our study, we overcame this problem. As mass loss effects did not differ significantly between *F. sylvatica* and *P. sylvestris*, we assume that coniferous tree species are more diverse at least in this relatively early successional stage. Note that in our study, *P. sylvestris* hosts on average approximately twice as many species as *F. sylvatica*, in terms of alpha diversity for rare and common species. However, more studies are needed which evaluate both diversity based on within-deadwood mycelia and the diversity of fruit bodies, standardized by decomposition over the course of succession, and considering tree species from different lineages.

Importantly, in both overall models for assessing the drivers of alpha and beta fungal diversity, the effect of the host tree species on diversity decreased between rare and dominant species. This may seem intuitive, but it has not often been explicitly tested and shown for fungal communities. A possible explanation for this finding could be that fungi that are abundant in deadwood are less host-dependent whereas more rare species are more dependent on the host tree species, perhaps due to habitat specificity - rare fungi are specialists of the host tree species (Crisfield et al., 2024). However, whether rare versus dominant species show higher degrees of specialization with their hosts and how community specialization is linked with species diversity patterns must be left to further studies.

It is notable that in the context of our experimental design involving canopy cover and deadwood enrichment, each of these two factors affected fungal diversity differently depending on the host tree species. Firstly, fungal alpha diversity in *F. sylvatica* deadwood was driven by canopy cover in that it was higher under closed canopies. It may be that, in the case of *F. sylvatica*, the conditions under the closed canopy (generally cooler and wetter) were conducive to fungal establishment and growth. Brabcová et al. (2022) and Perreault et al. (2023) also found that fungal community composition in wood-inhabiting fungi was driven in part by microclimate. Fungal beta diversity in *F. sylvatica* was also affected by canopy cover (but only for rare species). Secondly, the fungal alpha diversity of *P. sylvestris* was driven by deadwood resource availability, namely patches with habitat trees and tree crowns were linked with higher fungal alpha diversity and patches with crowns were linked to beta diversity (although only in the case of rare species). This effect in alpha diversity could be understood in the context of the “surface area effect” (Heilmann-Clausen & Christensen, 2004; Kruys & Jonsson, 1999). Assessing rarefaction curves, Heilmann-Clausen & Christensen (2004) found that small trees and branches hosted more fungal species per unit volume than larger trees and logs. This finding has been corroborated in Piché-Choquette et al. (2023). In Bässler et al. (2010), the authors also found higher fungal diversity in FWD than in CWD. In both studies, the authors hypothesized the cause for this as being the higher habitat diversity (per volume) in FWD. In FWD, there can be greater fluctuations in moisture and temperature and, for branches and tree crowns, there could be more extreme fluctuations in humidity and light intensity (Shiegg, 2001). Furthermore, habitat trees are also extremely heterogenous as a fungal resource. These structures offer a combination of healthy and damaged bark, sapwood, and tree crown, as well as the combination of CWD and FWD (i.e., branches and remaining parts of the tree crown). Perhaps it is this niche heterogeneity at the patch-level that contributes to the heterogeneous fungal communities found in *P. sylvestris*.

This finding overall supports our expectations to find higher fungal diversity in patches with deadwood enrichment compared to those where only stumps remain, due to the increased availability and heterogeneity of deadwood resources. However, the fact that environmental variables explaining fungal diversity differ strongly between tree species deserves further attention. Specifically, we cannot explain why the apparent niche heterogeneity effect observed for *P. sylvestris* was not visible for *F. sylvatica.* Additionally, the importance of canopy cover in explaining alpha diversity in *F. sylvatica* but not *P. sylvestris* is interesting. We speculate that this might be related to the basic structure of forest habitats; *F. sylvatica*- dominated forests are rather darker than *P. sylvestris*-dominated forests (as light transmission in forests correlates to the shade-tolerance, successional status, and foliage clumping of the dominant species, Lintunen et al., 2013). Such structural differences might have shaped the coevolutionary relationship between host tree species and their fungal inhabitants. In this vein, one might hypothesize that fungal species associated with *F. sylvatica*-dominated forests have evolved a stronger dependence on the relatively benign microclimate conditions of closed canopy forests than those species associated with *P. sylvestris*-dominated forests.

### 4.2 Effects of canopy, deadwood treatments, and host tree species on decomposition

The aim of the overall Structural Equation Model (SEM) was to test the direct and indirect (via diversity) effects of the canopy and deadwood enrichment treatments on deadwood mass loss. Based on this overall model one can assess the role of the host relative to the treatments. Such models are critical, for example, in the case that the host effect on fungal diversity (i.e., differences between *F. sylvatica* and *P. sylvestris*) might be confounded with differences in mass loss between tree species (Schreiber et al., 2025a). Nevertheless, the overall model together with the tree species specific models revealed interesting insights: First, the overall models, when considering all levels of diversity (i.e., alpha and beta across orders of *q*), revealed no consistent effect of the relationship between host tree species and mass loss.

Even though some models (i.e., SEMs which included beta diversity, PC2, across orders of *q*) showed that *F. sylvatica* was significantly more decomposed than *P. sylvestris*. Further, the effect, as can be seen in the raw data plots, was not very pronounced and seemed to be caused mainly by the higher (but not significant) level of decomposition of *F. sylvatica* under open canopies (Fig. 5). This contrasts with many studies showing higher decomposition rates in broad-leaf wood compared to coniferous wood (e.g., Kahl et al., 2017; Kipping et al., 2022). Differences in decomposition rates have been explained by wood traits like lignin and hemicellulose structure as well as nutrient content namely nitrogen (Weedon et al., 2009). And it has been shown that higher concentrations and a denser structure of lignin inhibits the decomposition of coniferous wood (e.g., Hatakka & Hammel, 2010). One reason why we observed only slight differences in mass loss between the host tree species might be that our study covered just the initial stages of decomposition.

Second, in the overall SEM, we revealed a significant negative relationship between mass loss and alpha diversity for common and dominant species. This negative effect might be partly explained by the lower mass loss and higher diversity of *P. sylvestris* compared to *F. sylvatica* (even though mass loss is not significantly different between tree species, see above). However, the effect sizes (i.e., *z* values) of the alpha diversity-decomposition relationships in the host tree species specific models were notably all negative, even though not significant in the full models (Table 3) but partly in the univariate models (Fig. 4). This overall indicates a negative diversity-mass loss relationship but we interpret our results with care as the critical values from the overall SEMs are not very large (Table S1). Together, our findings indicate antagonistic rather than synergistic mechanisms within the diversity-decomposition relationship. Our results support a lab experiment which, with a limited set of species, suggested that competitive interactions explain the negative diversity-decomposition relationship (Fukami et al., 2010). On the other hand, as mass loss (and decay) progress over time, thus reducing available fungal habitat, it may be that only a smaller subset of fungi are thus able to inhabit the substrate. However, whether the relationship between fungal diversity and decomposition is positive, neutral, or negative is far from clear. The available studies are inconsistent, which might be attributable to strong context dependencies (e.g., real-world, highly diverse communities versus lab experiments with selected species, as well as different environmental contexts and spatial scales – see Runnel et al., 2025 for a detailed discussion).

Third, overall SEM treatment effects may have been masked by contrasting host effects since the host tree species-specific models showed significant effects of the deadwood treatment while the overall model did not reveal any significant direct effects of the canopy and deadwood treatments. The results of the host tree species specific models on the effects of canopy cover and deadwood availability as well as alpha and beta diversity on wood mass loss in *F. sylvatica* and *P. sylvestris* deadwood indicate that canopy cover was not relevant to mass loss, contrary to our expectations (e.g., physical disintegration via radiation and photodegradation, see Introduction). Effects of the deadwood treatments were only significant and consistent across models for *F. sylvatica*. Mass loss of *F. sylvatica* deadwood increased in forest patches where trees had been totally removed and where crowns remained. Direct and independent effects from deadwood in the immediate surroundings on mass loss might be caused by modified microclimate effects. For example, a recent study demonstrated that deadwood temperature is lower in the summer months if the amount of deadwood in the surroundings is higher, which suggests buffering effects (Schreiber et al., 2025a). Thus, the positive effect of crowns on *F. sylvatica* deadwood mass loss might be caused by benign temperature and moisture allowing decomposer communities to increase their metabolic rate and thus decomposition capability (Schreiber et al., 2025b). However, it remains unclear why mass loss is then higher in patches with totally removed trees and why we did not observe similar effects with other types of deadwood enrichment. Here, more studies are needed to evaluate the physical conditions mediated by deadwood treatments that might modify metabolic rates of fungal communities adapted to different tree species.

## 5 Conclusions

Using a large-scale field experiment manipulating canopy cover, deadwood type and availability, we show that wood-inhabiting fungal diversity and deadwood decomposition are shaped primarily by host tree species identity rather than forest structure. This underscores the importance of maintaining a broad range of host species across forest landscapes to sustain fungal diversity. At the same time, species-specific management strategies are crucial: in *F. sylvatica* forests, maintaining heterogeneous canopy cover promotes complementary fungal communities, whereas in *P. sylvestris* forests, fungal diversity benefits from deadwood enrichment, including the retention of tree crowns after logging and the protection or creation of habitat trees through premature-senescence strategies. Because deadwood mass loss is driven by both deadwood structure and fungal diversity, decomposition processes are highly sensitive to forest management, but strongly dependent on dominant tree species. To better predict biodiversity and ecosystem functioning outcomes, future research should expand across spatial and temporal scales and develop mechanistic insights into fungal community assembly, coexistence and their functional roles in driving ecosystem processes.

## Notes

**Funding:** This work was supported by the Deutsche Forschungsgemeinschaft [BETA-FOR, 459717468].

### Competing Interest Statement

The authors have declared no competing interest.

