## Supplementary Material for "Experimentally manipulating forest structure to mimic management strategies: effects on deadwood fungal diversity and related ecosystem processes"

**Table S1. Piecewise Structural Equation Models (SEM)**

Summary of path significance resulting from the overall (i.e., including both *F. sylvatica* and *P. sylvestris* samples) Piecewise Structural Equation Models (SEM) across orders of *q*. For beta diversity, fungal community composition was represented by the first two axes of a Principal Coordinates Analysis (first axis: PC1; second axis: PC2). Critical values are given and significant group effects are indicated in bold. We log-transformed alpha diversity and square root transformed mass loss before modelling (see Materials and methods for more detail). For readability, only significant group effects are highlighted in bold. All significant effects are indicated with asterisks (* *p* < 0.05, ** *p* < 0.01, *** *p* < 0.001).

|  | *q* = 0, rare | | | *q* = 1, common | | | *q* = 2, dominant | |
| --- | --- | --- | --- | --- | --- | --- | --- | --- |
|  | **Crit. Value** | **P value** | | **Crit. Value** | **P value** | | **Crit. Value** | **P value** |
| Alpha Div. ~ | | | | | | | | |
| Canopy | 2.367 | | 0.124 | 2.171 | | 0.141 | 3.363 | 0.067 |
| Open | 43.211 | | <0.001*** | 23.098 | | <0.001*** | 18.227 | <0.001*** |
| Closed | 48.671 | | <0.001*** | 26.543 | | <0.001*** | 21.884 | <0.001*** |
| Deadwood | 4.999 | | 0.544 | 5.383 | | 0.496 | 8.502 | 0.204 |
| Stumps | 39.026 | | <0.001*** | 21.327 | | <0.001*** | 17.146 | <0.001*** |
| Trees removed | 25.789 | | <0.001*** | 14.563 | | 0.001** | 10.955 | 0.002** |
| Snags + Logs | 25.435 | | <0.001*** | 13.995 | | 0.001** | 10.198 | 0.002** |
| Habitat trees | 29.101 | | <0.001*** | 17.239 | | <0.001*** | 13.153 | 0.001** |
| Crowns | 26.120 | | <0.001*** | 15.776 | | 0.001** | 13.270 | 0.001** |
| Logs | 25.937 | | <0.001*** | 14.754 | | 0.001** | 11.029 | 0.002** |
| Snags | 25.794 | | <0.001*** | 14.982 | | 0.001** | 11.589 | 0.001** |
| Host tree | **93.432** | | **<0.001***** | **53.630** | | **<0.001***** | **22.582** | **<0.001***** |
| *F. sylvatica* | 42.397 | | <0.001*** | 21.999 | | <0.001*** | 18.194 | <0.001*** |
| *P. sylvestris* | 53.120 | | <0.001*** | 29.194 | | <0.001*** | 23.303 | <0.001*** |
| Marginal R^2^ | 0.50 | | | 0.37 | | | 0.26 | |
| Beta Div. (PC1) ~ | | | | | | | | |
| Canopy | 2.505 | | 0.114 | 2.730 | | 0.099 | 1.602 | 0.206 |
| Open | -1.485 | | 0.234 | -0.741 | | 0.513 | -0.707 | 0.530 |
| Closed | 0.636 | | 0.570 | 0.958 | | 0.409 | 0.529 | 0.633 |
| Deadwood | 10.222 | | 0.116 | 3.614 | | 0.729 | 4.592 | 0.597 |
| Stumps | 0.762 | | 0.501 | -0.883 | | 0.442 | -0.867 | 0.450 |
| Trees removed | 2.083 | | 0.129 | 0.553 | | 0.619 | -0.246 | 0.821 |
| Snags + Logs | -0.706 | | 0.531 | -0.018 | | 0.987 | -0.231 | 0.833 |
| Habitat trees | -0.919 | | 0.426 | 0.620 | | 0.579 | 0.410 | 0.709 |
| Crowns | -1.185 | | 0.321 | -0.611 | | 0.584 | -0.574 | 0.606 |
| Logs | -1.252 | | 0.299 | -0.387 | | 0.724 | -0.490 | 0.658 |
| Snags | -0.241 | | 0.826 | 0.865 | | 0.451 | 1.338 | 0.273 |
| Host tree | **1922.929** | | **<0.001***** | **2229.019** | | **<0.001***** | **1011.313** | **<0.001***** |
| *F. sylvatica* | -28.589 | | <0.001*** | -21.850 | | <0.001*** | -14.238 | 0.001** |
| *P. sylvestris* | 27.900 | | <0.001*** | 22.194 | | <0.001*** | 14.122 | 0.001** |
| Marginal R^2^ | 0.95 | | | 0.96 | | | 0.91 | |
| Beta Div. (PC2) ~ | | | | | | | | |
| Canopy | **4.391** | | **0.036*^1^** | 0.656 | | 0.418 | 1.789 | 0.181 |
| Open | 1.026 | | 0.380 | 0.934 | | 0.419 | -1.381 | 0.261 |
| Closed | -0.592 | | 0.596 | -0.077 | | 0.943 | 0.403 | 0.714 |
| Deadwood | 7.417 | | 0.284 | 2.996 | | 0.809 | 2.314 | 0.889 |
| Stumps | 0.770 | | 0.498 | 0.247 | | 0.821 | 0.011 | 0.992 |
| Trees removed | 1.136 | | 0.339 | 1.111 | | 0.348 | -0.749 | 0.508 |
| Snags + Logs | -0.651 | | 0.561 | -0.773 | | 0.496 | 0.669 | 0.552 |
| Habitat trees | -1.080 | | 0.359 | -0.017 | | 0.987 | -0.544 | 0.624 |
| Crowns | -0.199 | | 0.855 | -0.172 | | 0.874 | -0.052 | 0.962 |
| Logs | 0.777 | | 0.494 | 1.018 | | 0.384 | -1.129 | 0.341 |
| Snags | 0.587 | | 0.598 | 0.363 | | 0.741 | -0.120 | 0.912 |
| Host tree | 0.007 | | 0.935 | 0.062 | | 0.804 | 0.390 | 0.532 |
| *F. sylvatica* | 0.273 | | 0.802 | 0.317 | | 0.772 | -0.145 | 0.894 |
| *P. sylvestris* | 0.229 | | 0.834 | 0.623 | | 0.578 | -0.956 | 0.410 |
| Marginal R^2^ | 0.11 | | | 0.04 | | | 0.06 | |
| Mass loss ~ | | | | | | | | |
| Canopy | 0.002 | | 0.965 | 0.002 | | 0.965 | 0.017 | 0.896 |
| Open | 15.990 | | <0.001*** | 16.283 | | 0.001** | 16.241 | 0.001** |
| Closed | 17.355 | | <0.001*** | 17.730 | | <0.001*** | 17.800 | <0.001*** |
| Deadwood | 8.551 | | 0.201 | 8.210 | | 0.223 | 9.635 | 0.141 |
| Stumps | 12.719 | | 0.001** | 12.971 | | 0.001** | 12.909 | 0.001** |
| Trees removed | 9.523 | | 0.003** | 9.577 | | 0.002** | 9.567 | 0.002** |
| Snags + Logs | 9.122 | | 0.003** | 9.051 | | 0.003** | 8.957 | 0.003** |
| Habitat trees | 7.500 | | 0.005** | 7.749 | | 0.005** | 7.691 | 0.005** |
| Crowns | 9.442 | | 0.003** | 9.742 | | 0.002** | 9.914 | 0.002** |
| Logs | 8.167 | | 0.004** | 8.312 | | 0.004** | 8.280 | 0.004** |
| Snags | 8.174 | | 0.004** | 8.445 | | 0.004** | 8.525 | 0.003** |
| Host tree | 0.143 | | 0.705 | 0.054 | | 0.816 | 0.659 | 0.417 |
| *F. sylvatica* | 14.609 | | <0.001*** | 15.924 | | 0.001** | 17.399 | <0.001*** |
| *P. sylvestris* | 14.977 | | <0.001*** | 16.324 | | 0.001** | 16.837 | 0.001** |
| Alpha div. | -1.563 | | 0.126 | **-2.440** | | **0.019*** | **-2.499** | **0.017*** |
| Marginal R^2^ | 0.15 | | | 0.18 | | | 0.18 | |
| Mass loss ~ | | | | | | | | |
| Canopy | 0.143 | | 0.705 | 0.045 | | 0.833 | 0.056 | 0.814 |
| Open | 16.177 | | 0.001** | 16.067 | | 0.001** | 16.096 | 0.001** |
| Closed | 16.979 | | <0.001*** | 16.934 | | <0.001*** | 17.000 | <0.001*** |
| Deadwood | 9.129 | | 0.166 | 9.223 | | 0.161 | 9.114 | 0.167 |
| Stumps | 12.377 | | 0.001** | 12.453 | | 0.001** | 12.465 | 0.001** |
| Trees removed | 9.092 | | 0.003** | 9.447 | | 0.003** | 9.428 | 0.003** |
| Snags + Logs | 9.137 | | 0.003** | 9.111 | | 0.003** | 9.101 | 0.003** |
| Habitat trees | 7.313 | | 0.005* | 7.300 | | 0.005** | 7.296 | 0.005** |
| Crowns | 9.320 | | 0.003** | 9.272 | | 0.003** | 9.282 | 0.003** |
| Logs | 8.342 | | 0.004** | 8.293 | | 0.004** | 8.287 | 0.004** |
| Snags | 8.384 | | 0.004** | 8.376 | | 0.004** | 8.336 | 0.004** |
| Host tree | 0.993 | | 0.319 | 0.002 | | 0.965 | 0.095 | 0.758 |
| *F. sylvatica* | 6.432 | | 0.008** | 5.403 | | 0.012* | 7.815 | 0.004** |
| *P. sylvestris* | 4.706 | | 0.018* | 5.299 | | 0.013* | 7.308 | 0.005** |
| Beta div. (PC1) | 0.523 | | 0.604 | -0.384 | | 0.703 | -0.329 | 0.744 |
| Marginal R^2^ | 0.13 | | | 0.13 | | | 0.13 | |
| Mass loss ~ | | | | | | | | |
| Canopy | 0.025 | | 0.875 | 0.032 | | 0.858 | 0.005 | 0.942 |
| Open | 16.230 | | 0.001** | 16.262 | | 0.001** | 16.136 | 0.001** |
| Closed | 17.054 | | <0.001*** | 17.289 | | <0.001*** | 17.372 | <0.001*** |
| Deadwood | 9.495 | | 0.148 | 9.457 | | 0.150 | 9.815 | 0.133 |
| Stumps | 12.027 | | 0.001** | 12.603 | | 0.001** | 12.670 | 0.001** |
| Trees removed | 9.374 | | 0.003** | 9.338 | | 0.003** | 9.435 | 0.003** |
| Snags + Logs | 9.172 | | 0.003** | 9.297 | | 0.003** | 9.336 | 0.003** |
| Habitat trees | 7.325 | | 0.005** | 7.388 | | 0.005** | 7.306 | 0.005** |
| Crowns | 9.383 | | 0.003** | 9.445 | | 0.003** | 9.447 | 0.003** |
| Logs | 8.321 | | 0.004** | 8.258 | | 0.004** | 8.224 | 0.004** |
| Snags | 8.393 | | 0.004** | 8.425 | | 0.004** | 8.480 | 0.003** |
| Host tree | **4.298** | | **0.038*** | **4.527** | | **0.033*** | **4.846** | **0.028*** |
| *F. sylvatica* | 18.806 | | <0.001*** | 18.904 | | <0.001*** | 18.982 | <0.001*** |
| *P. sylvestris* | 16.322 | | 0.001** | 16.344 | | 0.001** | 16.284 | 0.001** |
| Beta div. (PC2) | 0.870 | | 0.389 | 1.387 | | 0.173 | -1.595 | 0.118 |
| Marginal R^2^ | 0.13 | | | 0.14 | | | 0.15 | |

^1^Although the SEM built-in Anova indicated the group effect was significant, the post hoc follow-up test yielded no significant results. Therefore, for our purposes, we do not consider this result as significant.

**Table S2.** **Host tree species specific beta regression models**

Test statistics and *p*-values from fitting a beta regression model (*gam* function, *mgcv* package) on mass loss in response to canopy and deadwood treatments and for each order of *q* and for each host tree species separately. The reference level for the canopy treatment is stated in parentheses. The reference level for the deadwood treatment was stumps, i.e., patches where only stumps were left remaining. Significant effects are indicated with asterisks (* *p* < 0.05, ** *p* < 0.01, *** *p* < 0.001).

|  | Fixed effects | *q* = 0, rare | | *q* = 1, common | | *q* = 2, dominant | |
| --- | --- | --- | --- | --- | --- | --- | --- |
|  |  | **z value** | **Pr(>\|z\|)** | **z value** | **Pr(>\|z\|)** | **z value** | **Pr(>\|z\|)** |
| *F. sylvatica* | Canopy (Open) | -0.094 | 0.925 | -0.004 | 0.997 | 0.069 | 0.945 |
|  | Trees removed | **2.559** | **0.011*** | **2.445** | **0.014*** | **2.376** | **0.018*** |
|  | Snags + Logs | 1.749 | 0.080 | 1.656 | 0.098 | 1.532 | 0.126 |
|  | Habitat trees | 0.906 | 0.365 | 0.788 | 0.431 | 0.592 | 0.554 |
|  | Crowns | **2.900** | **0.004**** | **2.794** | **0.005**** | **2.859** | **0.004**** |
|  | Logs | 1.506 | 0.132 | 1.487 | 0.137 | 1.403 | 0.160 |
|  | Snags | 0.610 | 0.542 | 0.574 | 0.566 | 0.461 | 0.645 |
|  | Alpha diversity | -0.391 | 0.696 | -0.935 | 0.350 | -1.385 | 0.166 |
|  | **Deviance explained** | 27.7% | | 38.4% | | 28.7% | |
|  | Canopy (Open) | -0.319 | 0.750 | -0.215 | 0.830 | -0.231 | 0.817 |
|  | Trees removed | **2.190** | **0.028*** | **2.470** | **0.014*** | **2.479** | **0.013*** |
|  | Snags + Logs | 1.701 | 0.089 | 1.716 | 0.086 | 1.677 | 0.093 |
|  | Habitat trees | 0.817 | 0.414 | 0.813 | 0.416 | 0.799 | 0.424 |
|  | Crowns | **2.964** | **0.003**** | **2.929** | **0.003**** | **2.913** | **0.004**** |
|  | Logs | 1.645 | 0.100 | 1.555 | 0.120 | 1.544 | 0.123 |
|  | Snags | 0.571 | 0.568 | 0.549 | 0.583 | 0.476 | 0.634 |
|  | Beta diversity (PC1) | 0.880 | 0.379 | 0.343 | 0.731 | 0.441 | 0.659 |
|  | **Deviance explained** | 25.3% | | 27.7% | | 28.0% | |
|  | Canopy (Open) | -0.167 | 0.867 | -0.211 | 0.833 | -0.162 | 0.871 |
|  | Trees removed | **2.578** | **0.010*** | **2.605** | **0.009**** | **2.550** | **0.011*** |
|  | Snags + Logs | 1.642 | 0.101 | 1.592 | 0.111 | 1.731 | 0.084 |
|  | Habitat trees | 0.845 | 0.398 | 0.887 | 0.375 | 0.911 | 0.362 |
|  | Crowns | **2.856** | **0.004**** | **2.899** | **0.004**** | **2.920** | **0.004**** |
|  | Logs | 1.564 | 0.118 | 1.595 | 0.111 | 1.546 | 0.122 |
|  | Snags | 0.666 | 0.506 | 0.704 | 0.481 | 0.661 | 0.509 |
|  | Beta diversity (PC2) | -0.167 | 0.867 | -0.497 | 0.619 | 0.075 | 0.940 |
|  | **Deviance explained** | 26.7% | | 26.5% | | 26.8% | |
| *P. sylvestris* | Canopy (Open) | 0.500 | 0.617 | 0.526 | 0.599 | 0.734 | 0.463 |
|  | Trees removed | 0.528 | 0.598 | 0.544 | 0.587 | 0.523 | 0.601 |
|  | Snags + Logs | 0.986 | 0.324 | 0.713 | 0.476 | 0.602 | 0.547 |
|  | Habitat trees | -1.754 | 0.079 | -1.406 | 0.160 | -1.312 | 0.190 |
|  | Crowns | -0.276 | 0.782 | 0.123 | 0.902 | 0.262 | 0.793 |
|  | Logs | -0.411 | 0.681 | -0.564 | 0.573 | -0.489 | 0.625 |
|  | Snags | -0.294 | 0.769 | -0.163 | 0.871 | 0.131 | 0.896 |
|  | Alpha diversity | -1.091 | 0.275 | -1.648 | 0.099 | -1.491 | 0.136 |
|  | **Deviance explained** | 35.9% | | 38.8% | | 39.8% | |
|  | Canopy (Open) | 0.464 | 0.642 | 0.555 | 0.579 | 0.574 | 0.566 |
|  | Trees removed | 0.471 | 0.637 | 0.616 | 0.538 | 0.472 | 0.637 |
|  | Snags + Logs | 1.282 | 0.200 | 1.100 | 0.271 | 0.959 | 0.338 |
|  | Habitat trees | -1.069 | 0.285 | -1.696 | 0.090 | -1.745 | 0.081 |
|  | Crowns | 0.154 | 0.877 | -0.318 | 0.750 | -0.352 | 0.725 |
|  | Logs | -0.030 | 0.976 | -0.208 | 0.835 | -0.256 | 0.798 |
|  | Snags | 0.885 | 0.376 | 0.229 | 0.819 | 0.259 | 0.796 |
|  | Beta diversity (PC1) | 1.411 | 0.158 | -0.521 | 0.602 | -0.802 | 0.422 |
|  | **Deviance explained** | 39.9% | | 37.0% | | 38.1% | |
|  | Canopy (Open) | 0.751 | 0.453 | 0.003 | 0.997 | 0.325 | 0.745 |
|  | Trees removed | 0.485 | 0.628 | 0.745 | 0.456 | 0.967 | 0.334 |
|  | Snags + Logs | 1.414 | 0.157 | 1.032 | 0.302 | 1.575 | 0.115 |
|  | Habitat trees | -1.039 | 0.299 | -1.559 | 0.119 | -1.382 | 0.167 |
|  | Crowns | -0.081 | 0.936 | 0.325 | 0.745 | 0.383 | 0.702 |
|  | Logs | -0.131 | 0.896 | -0.087 | 0.931 | 0.039 | 0.969 |
|  | Snags | 0.395 | 0.693 | 0.560 | 0.575 | 0.515 | 0.607 |
|  | Beta diversity (PC2) | 1.527 | 0.127 | **2.588** | **0.010*** | **-2.381** | **0.017*** |
|  | **Deviance explained** | 40.2% | | 43.8% | | 42.4% | |
